# Flies getting filthy: The precopulatory mating behaviours of three mud-dwelling species of Australian *Lispe* (Diptera: Muscidae)

**DOI:** 10.1101/2021.04.22.439303

**Authors:** Nathan J. Butterworth, James F. Wallman

## Abstract

*Lispe* (Diptera: Muscidae) is a cosmopolitan genus of predatory flies that inhabit the muddy and sandy surrounds of water bodies. There are more than 163 described species worldwide, many of which are known to exhibit cursorial courtship displays which involve complex visual and vibratory signals. Despite the widespread distribution of these flies and their remarkable courtship displays, the biology and behaviour of most species are entirely unknown. Here, for the first time, we describe the pre-copulatory mating behaviours of three widespread and common Australian species: *Lispe sydneyensis, Lispe albimaculata* and *Lispe xenochaeta*. We demonstrate that all three species exhibit entirely unique courtship displays, consisting of complex behavioural repertoires. Importantly, we highlight intrasexual competition in *L. sydneyensis*, where males engage in competitive dances and combat. We also report female-male aggression in *L. albimaculata* and *L. xenochaeta* where females charge and display towards males. These novel mating systems provide unique opportunities to test ecological and evolutionary hypotheses.

## INTRODUCTION

The dipteran clade Calyptratae is incredibly diverse with more than 18,000 described species, including some of the most well-known flies such as house flies, blow flies, flesh flies, and bot flies (Kutty et al. 2010). These flies express an astounding variety of complex sexual behaviours including the sensual dances of the waltzing blowfly *Chrysomya flavifrons* (Calliphoridae) (Butterworth et al. 2019), the high-speed courtship flights of the lesser house fly *Fannia canicularis* (Muscidae) (Land and Collett 1974), and the flashy mating displays of the satellite fly *Phrosinella aurifacies* (Sarcophagidae) (Spofford and Kurczewski 1985). However, one particular genus of muscid flies – *Lispe* – has taken these sexual innovations to the extreme.

*Lispe* is a cosmopolitan genus of flies which inhabit open sandy or muddy substrates surrounding puddles, creeks, rivers, lakes, and beaches (Werner and Pont 2006; Zhang et al. 2013; Fogaça and de Carvalho 2018). The group is characterised by the enlarged facial palps, which have been adapted for sexual signalling in some species (White et al. 2020a). There are more than 163 species worldwide (Pont 2019) all of which appear to be predators and scavengers of small invertebrates or their remains (Werner and Pont 2006; Vikhrev 2011). Most species seem to exhibit unique and complex courtship displays, such as the circular cavorting of *Lispe tentaculata* along the muddy banks of rivers in Europe (Frantsevich and Gorb 2006), or the iridescent face-to-face dances of *Lispe cana* along the coastal beaches of Australia (White et al. 2020a; White et al. 2020b). Australia is home to at least 39 species of *Lispe* (Pont 2019), which due to their widespread abundance and diverse behaviours have exceptional potential as models for testing ecological and evolutionary hypotheses (White et al. 2020b).

Despite their unique ecologies and mating systems, the biology of almost all Australian *Lispe* species (besides *L. cana*) remains entirely unknown (Pont 2019). Here, for the first time, we report the diverse courtship behaviours of three mud-dwelling Australian species: *Lispe sydneyensis, Lispe albimaculata* and *Lispe xenochaeta*. These species provide unique opportunities for field studies of evolution and behaviour because they are common and broadly distributed throughout Australia, exhibit remarkably diverse mating systems, and are easy to observe and film.

## METHODS

### Field site

Filming was conducted between the 6^th^ and 12^th^ of September 2020 around muddy pools along a sandy track in Huskisson, NSW Australia (35°02’59.0”S 150°40’07.4”E) (Figure 1). All observations were made between 10:00 and 15:00 under natural light and temperature conditions (temperature min: 17.8°C, max: 21.0°C, mean: 19.4°C). The last period of substantial rain (daily amounts exceeding 10 mm) was between the 8^th^ and 10^th^ August, and as such the bodies of water must have been present for several weeks prior to observation and were inundated with frog and mosquito larvae.

**Figure 1.**
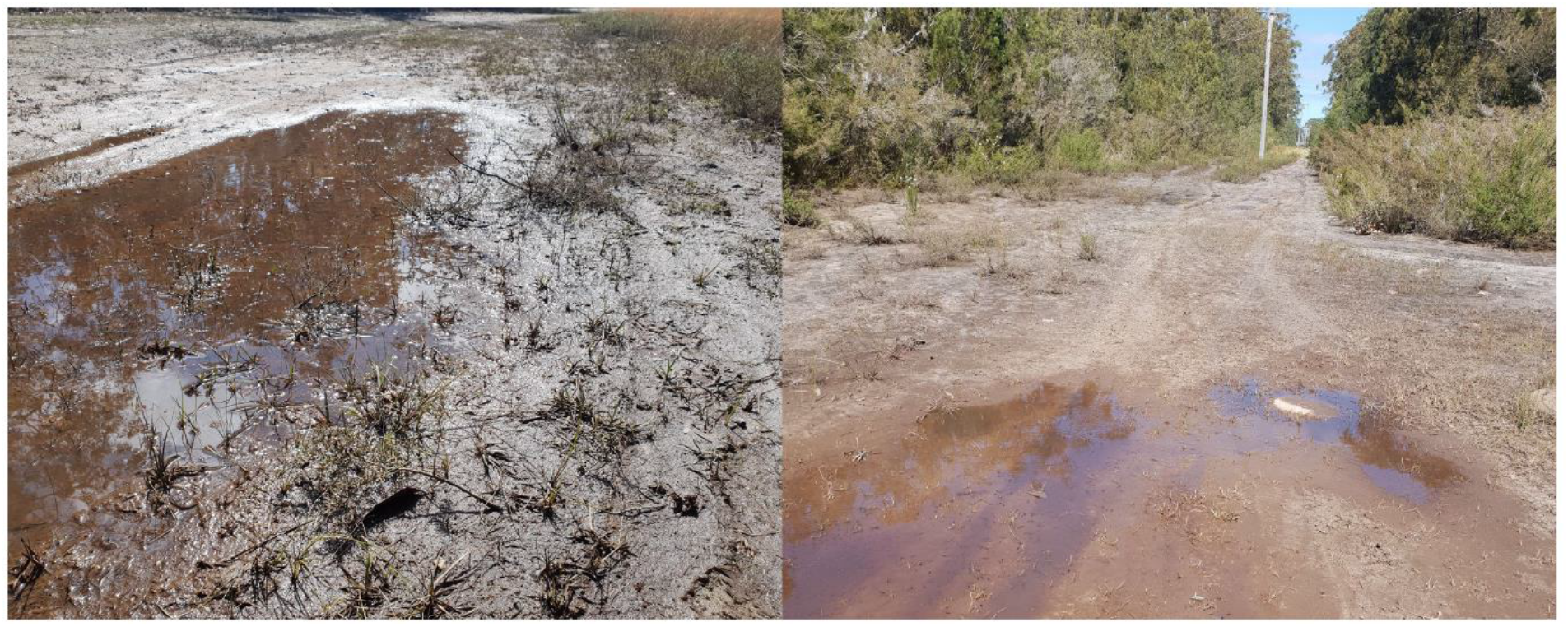
Muddy site where observations were made. Water had been present for approximately 2-3 weeks and was inundated with frog and mosquito larvae. Both *L. sydneyensis* and *L. albimaculata* were in high abundance (∼50-100 individuals at any time), while *L. xenochaeta* were less frequently observed (only 1-2 individuals at any time).

### Insect identification

To identify species and correctly assign courtship behaviours, for each species between two and four courting pairs were captured and euthanised. Taxonomic identification followed the taxonomic key of Pont (2019) alongside comparison with museum specimens from the Australian National Insect Collection (ANIC). Three species were identified (Figure 2).

**Figure 2.**
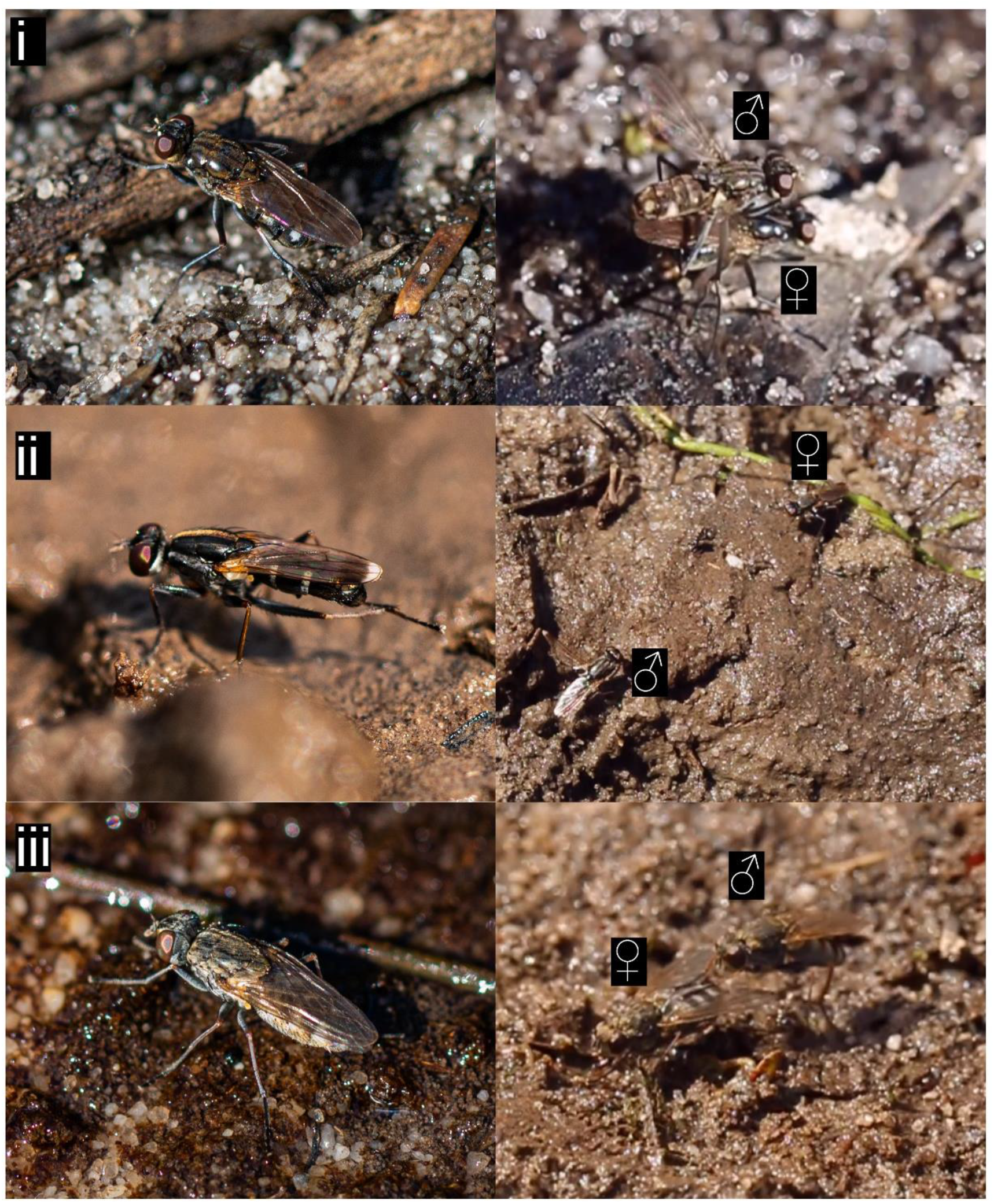
The three species of Australian *Lispe* found courting around the muddy pool. Pictured are (i) *L. sydneyensis* and a male ‘straddling’ a female, (ii) *L. albimaculata* and a male ‘wing-revealing’ towards a female, and (iii) *L. xenochaeta* and a male ‘holding’ a female. Photos were taken with a Canon 70D DSLR camera with a Canon EF 100mm f/2.8L lens. Photos credited to Nathan Butterworth.

### Behavioural observations

Conspecific interactions (male-female, female-male, and male-male) were recorded with a Canon 70D DSLR camera with a Canon EF 100mm f/2.8L lens. Filming continued until one or both flies left the area and could no longer be observed. Once video footage was obtained, slow-motion playback with Adobe Premiere Pro allowed us to describe all inter-and intra-sexual interactions.

## RESULTS

A total of 57 individual interactions were recorded across the three species. For *L. sydneyensis* we recorded thirty-nine interactions, for *L. albimaculata* we recorded sixteen interactions, and for *L. xenochaeta* we recorded two interactions. From this footage, we were able to describe the behaviours expressed by these species during courtship. For video footage of each of the behaviours, refer to Supplementary Materials 1-3.

### The straddling mud fly, *Lispe sydneyensis*

This was the most common species at the site. Notably, *L. sydneyensis* males have greatly elongated mid-legs (Figure 3.i), which allows them to position their entire body atop the back of the female and to remain in this position as the female moves around (Fig 2.i). The species also has iridescent markings on the head and palps which may be involved in courtship (Figure 3.ii). The male courtship display is complex, involving several discrete behaviours (Table 1; Supplementary Material 1). In sequential order, the male ‘orients’ towards the female, until he is within ∼5 mm, at which point he rapidly encircles her while waving his mid-legs. He then makes a few ‘straddle-strikes’ onto the back of the female, before committing to the final straddling position. While ‘straddling’, the male vigorously vibrates his wings and strokes the head and wings of the female with his fore-and hind-legs, respectively. Females seem unaffected by these behaviours, and continue to explore, preen themselves, and feed on surrounding matter. After a certain period of ‘straddling’, the male attempts copulation with the female, although we only observed this on one occasion. We did not observe any female-specific mating behaviours or responses to male mating attempts. However, there is clearly intense male-male competition for females. We observed numerous encounters between males, where they approach each other while rapidly moving their bodies up and down (‘bopping’), and in some cases waving their mid-legs. If neither male concedes, this often leads to a frontal attack with the proboscis, usually resulting in a brief tug-of-war between the two.

**Figure 3.**
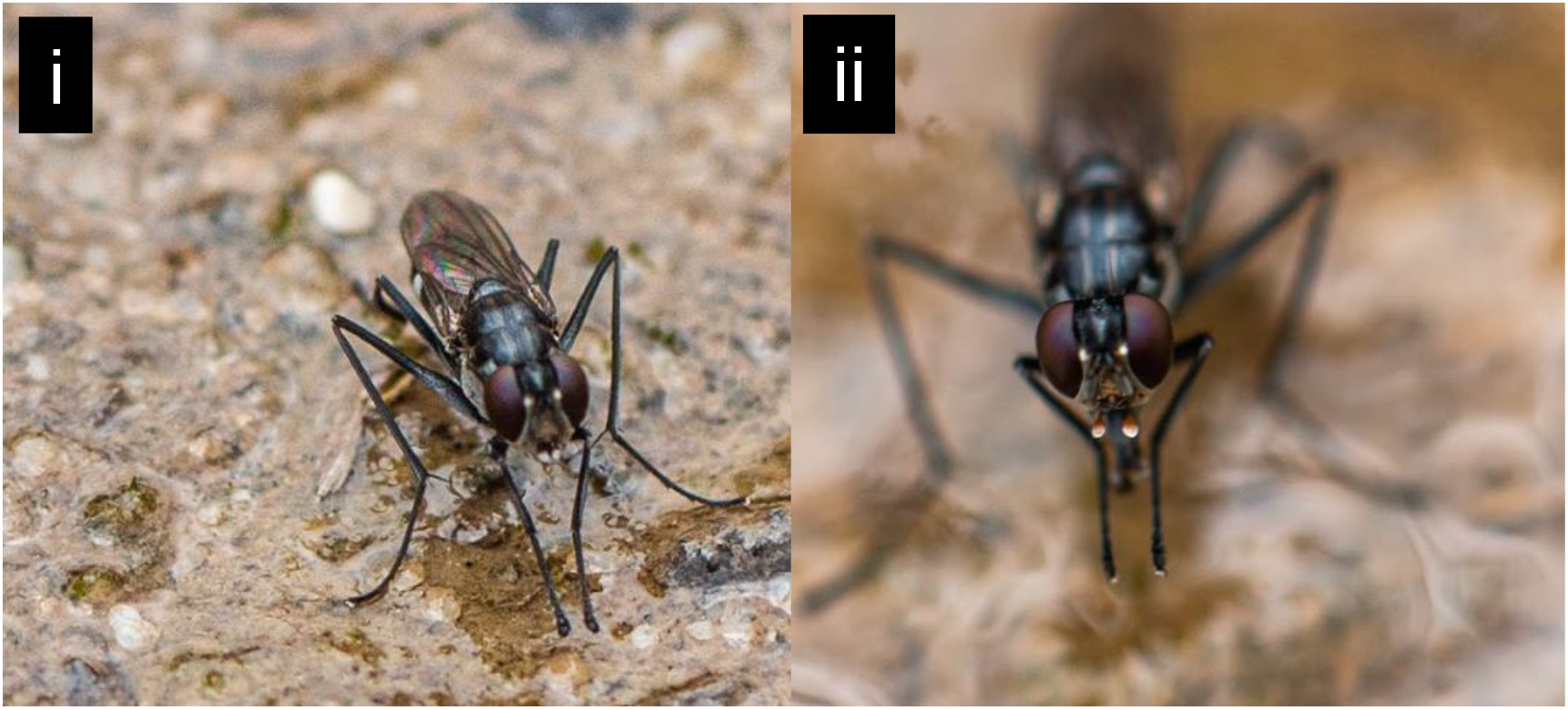
**i)** Elongated mid-legs and **ii)** iridescent spots seen on the head and facial palps of *Lispe sydneyensis*. The elongated mid-legs allow the male to position his entire body atop the back of the female (Figure 2.i). The iridescent spots may play an important role in sexual signalling, like the iridescent facial features involved in the sexual behaviour of *Lispe cana* (White et al. 2019). Photos were taken with a Canon 70D DSLR camera with a Canon EF 100mm f/2.8L lens. Photos credited to Nathan Butterworth.

**Table 1.** An ethogram describing the sexual behaviours displayed by male and female *L. sydneyensis*. All behaviours were observed in at least three of the recorded observations (Total N = 39). ‘M-F’ represents behaviours directed from a male to a female, ‘M-M from a male to a male, and ‘F-M’ from a female to a male.

| Direction | Behaviour | Description |
| --- | --- | --- |
| M-F | Orienting | Upon encountering a female, the male follows and attempts to position himself near her (within ~5 mm). |
|  | Circling | The male rapidly encircles the female, usually in conjunction with mid-leg waving. |
|  | Mid-leg waving | Immediately prior to straddling the female, the male rapidly waves both of his mid-legs. |
|  | Straddle-strike | The male quickly jumps on and off the back of the female, often several times immediately prior to straddling. |
|  | Straddling | The male sits atop the female, with a mid-leg on either side, which stabilises him while the female walks around. |
|  | Wing-vibrating | The male rapidly vibrates both of his wings while straddling the female. |
|  | Stroking | While straddling the female, the male strokes her head and wings by shaking his fore-legs and hind-legs, respectively. |
| M-M | Bopping | When two males encounter each other, they use their legs to repeatedly raise and then lower their bodies as they slowly approach each other. |
|  | Fighting | If neither male concedes during bopping, then one male initiates a frontal attack with the proboscis, sometimes resulting in a tug-of-war between the two. |
|  | Mid-leg waving | The male rapidly waves his mid legs while facing towards a competing male. |
| F-M | None observed* | — |

\*There may be behaviours in this category; more observations are required.

### The matador mud fly, *Lispe albimaculata*

This species was also commonly encountered. Both males and females exhibit white tips to their wings, which seems to play an integral role in the male display (Figure 2.ii). The courtship consists of a complex back and forth between the sexes, with both the male and the female exhibiting several discrete behaviours (Table 2; Supplementary Material 2). In sequential order, the male ‘orients’ towards the female, until within ∼10 mm of her. The male then ‘sneaks’ slowly closer to her and then remains perfectly still for a prolonged period (anywhere from a few seconds to more than one minute), until the female rapidly spins to ‘face-off’ with the male. He then flicks out one of his white-tipped wings and rapidly vibrates it (although vibration is not always involved). This usually results in an immediate response by the female, who will face the male and move towards the wing to inspect it. At this point, the male either immediately attempts to force copulation or remains stationary for a period (sometimes for several minutes) before doing so. In response to this copulation attempt, females often become aggressive, resulting in a tussle between the two. We only observed one interaction where the female eventually relented and accepted mating by the male. In certain cases where the females are entirely non-receptive, they will lower their heads, spread their wings, and sometimes stomp their mid-legs – which appears to deter males in most circumstances. We observed no competitions or behavioural interactions between males.

**Table 2.** An ethogram describing the sexual behaviours displayed by male and female *L. albimaculata*. All behaviours were observed in at least three of the recorded observations (Total N = 16). ‘M-F’ represents behaviours directed from a male to a female, ‘M-M from a male to a male, and ‘F-M’ from a female to a male.

| Direction | Behaviour | Description |
| --- | --- | --- |
| M-F | Orienting | Upon encountering a female, the male moves towards her until within ~10 mm. |
|  | Sneak | Once within ~10 mm of the female, the male directly faces one side of her body, and inches forward at a slow pace, eventually remaining completely stationary. |
|  | Wing reveal | Within ~5 mm of the female, and when completely stationary, the male quickly flicks out one wing (left or right) and in some cases rapidly vibrates it. |
|  | Tackle | Once noticed by the female, the male swiftly jumps towards the female, presumably to force copulation. |
| M-M | None observed* | – |
| F-M | Wing spread | In response to a male’s courtship attempt, the female positions her head towards the ground, spreading her wings while facing directly towards the male. |
|  | Face-off | In response to the presence of a sneaking male, the female rapidly spins to face the male, then remains stationary for several seconds. |
|  | Approach | Following the face-off, the female slowly makes a frontal approach towards the male’s wing. |
|  | Wing inspection | The female sits directly next to the male’s revealed wing for several milliseconds, before turning around. |
|  | Mid-leg thumping | The female stomps both mid-legs while facing an approaching male. |
|  | Charge | Upon encountering a male, the female makes an aggressive charge towards him. |

\*There may be behaviours in this category; more observations are required.

### The hopping mud fly, *Lispe xenochaeta*

This species was only seen courting twice over the four days of filming. This was by far the most difficult species to film, as there were numerous complex interactions between the sexes (Table 3; Supplementary Material 3) and females would move frantically around the environment. In sequential order, the male ‘orients’ towards the moving female, closely following her before performing a series of sideways ‘hops’ (after each hop returning to his initial position) occasionally followed by attempts to ‘tackle’ her. In response, the female sometimes makes a frontal ‘charge’ towards the male. If the female is receptive, she continues to move around the environment while ‘quivering’ and ‘spreading’ her wings to reveal her black and white patterned abdomen. After several minutes of this back and forth, the receptive female comes to a standstill at which point the male proceeds to ‘hold’ her abdomen (Fig 2.iii) and begins ‘thumping’ his mid-legs and occasionally flicking his wings. After a period (between 30 seconds and several minutes) of ‘thumping’ and ‘wing-flicking’, the male attempts copulation. If the female is not receptive, she frantically shakes her body until the male detaches. We observed no interactions between males.

**Table 3.** An ethogram describing the sexual behaviours displayed by male and female *L. xenochaeta*. All behaviours were observed in both recorded observations (Total N = 2). ‘M-F’ represents behaviours directed from a male to a female, ‘M-M from a male to a male, and ‘F-M’ from a female to a male.

| Direction | Behaviour | Description |
| --- | --- | --- |
| M-F | Orienting | Upon encountering a female, the male pursues and attempts to orient himself behind her. |
|  | Wing extension | While orienting and facing the female, the male extends both wings. |
|  | Hopping | Facing the female, the male performs several sideways hops, each time returning to his starting position before hopping again. |
|  | Tackle | The male swiftly flies or jumps toward the female, striking her with his body. This may be a preliminary attempt at copulation. |
|  | Holding | The male positions himself behind the female and holds her abdomen or wings with his forelegs. |
|  | Thumping | While holding the female, the male vigorously thumps his mid-legs onto the ground. |
|  | Wing flicking | While holding and thumping, the male occasionally flicks both wings. |
| M-M | None observed* | — |
| F-M | Wing spread | When pursued by a male, the female continues to move while spreading both wings to reveal her patterned abdomen. |
|  | Wing quiver | The female will rapidly quiver both wings, often while they are spread open. |
|  | Charge | When being courted by a male, the female makes an aggressive charge directly towards the front of the male. |

\*There may be behaviours in this category; more observations are required.

## DISCUSSION

The genus *Lispe* is found in every biogeographic region (Pont 2019) and most described species seem to exhibit some form of cursorial courtship (Werner and Pont 2006). These courtship displays consist of species-specific behavioural repertoires (Frantsevich and Gorb 2006; Werner and Pont 2006; White et al. 2020) which suggests that sexual selection has played an important role in the evolution of the genus. Here, for the first time, we describe the pre-copulatory sexual behaviour of three Australian species: *L. sydneyensis, L. albimaculata* and *L. xenochaeta*. Considering that these species are common and easy to observe around ephemeral pools, they provide promising opportunities for future studies of evolution and behaviour.

### The straddling mud fly, *Lispe sydneyensis*

The most striking feature of this species is the elongation of the mid-legs in males and the associated straddling behaviour. The ‘straddling’ behaviour of *L. sydneyensis* is similar to the ‘holding’ in *L. xenochaeta* and ‘straddling’ seen in *L. cana* (White et al. 2019) – suggesting that straddling behaviours are widespread in *Lispe*. In the latter two species, males hold on to the back of the female with their forelegs, follow the female around, and only mount during copulation. The key difference in *L. sydneyensis* is that the male sits entirely atop the female and balances himself with the mid-legs while following her movements. Broadly, these ‘holding/straddling’ behaviours probably select for the ability of males to closely follow females and to guard her from nearby competitors prior to copulation. Such forms of precopulatory mate guarding are seen in many other insects including saproxylic parasitoid wasps (Hymenoptera: Ibaliidae) (Kuramitsu et al. 2019) and black scavenger flies (Diptera: Sepsidae) (Pont and Meier 2002). Pre-copulatory mate guarding usually evolves in response to high levels of male-male competition, which may result from a male-biased sex ratio – as appears to be the case in both *L. sydneyensis* and *L. cana* (personal observation). However, it is also plausible that the male ‘straddling’ seen in *L. sydneyensis* is used to access the female’s viewpoint, as is the case in *L. cana* (White et al. 2020a). By aligning his field-of-view with the female’s, the male can determine when the female is viewing a background against which he will stand out, allowing him to time maximise the salience of his display. In support of this, there are iridescent spots on the head and facial palps of *L. sydneyensis*, which are only visible at certain angles (Figure 3i) and may act as visual signals, akin to the facial colouration seen in *Lispe cana* (White et al. 2020a). Notably, the males vigorously vibrate their wings for the entire period they are atop the females. It is likely that this energetically costly performance produces aural cues, as in many *Drosophila* (Morley et al. 2012). The duration for which a male can remain atop the female as well as vibrate his wings may also act as an honest signal of male quality. Lastly, we observed high levels of male-male competition whereby males would frequently engage in one-on-one ‘bopping’ which often led to fights. Male-male ‘bopping’ seems to allow males to assess the quality of their competitors before fighting – as not all instances of ‘bopping’ led to fighting – which suggests that males adjust their tactics according to their rival’s quality (Swierk and Langkilde 2013). Male-male competition is widespread in flies. Other notable examples include mushroom flies of the genus *Tapeigaster* (Diptera: Heleomyzidae) (McAlpine and Kent 1981) and antler flies of the genus *Protopiophila* (Diptera: Piophilidae). However, *L. sydneyensis* makes a particularly good system for investigating the intricacies of male-male competition, because the species is easy to find, film, collect in large numbers, and male-male encounters are frequent.

### The matador mud fly, *Lispe albimaculata*

This species is unique in both sexes having white tips to their wings, which the males display as they ‘wing-reveal’ during courtship. This white wing tip is probably a species-specific signal, as is seen in many other insects (Fordyce et al. 2002; Butterworth et al. 2019; Butterworth et al. 2021). The vibrations that the males exhibit during wing-reveal may be associated with acoustic cues similar to many other fly species (Benelli et al. 2012). Notably, female-male aggression is common during courtship, whereby females are often seen attacking or ‘charging’ towards males. Females also exhibit ‘wing-spread’ and ‘mid-leg thumping’ when males approach, which appear to be signals of rejection, similar to the wing-vibrations used by females of the yellow dung fly *Scatophaga stercoraria* to signal non-receptivity (Parker 1970). The use of the mid-legs during courtship seems to be an ancestral trait that has been adapted for various purposes in *Lispe*, such as ‘thumping’ in *L. xenochaeta* and ‘mid-leg waving’ in *Lispe sydneyensis*. Regarding the aggressive behaviours, males of *L. albimaculata* often try to force copulation, so it may be that female aggression evolved in response to male aggression (Arnqvist and Henriksson 1997; Hohmann and Fruth 2003; Maklakov et al. 2004). It is also plausible that female aggression occurs post-mating after receipt of a male’s ejaculate and a subsequent reduction in sexual receptivity, as in *Drosophila* (Bath et al. 2017; Bath et al. 2021), or that aggression is related to the increased risk of predator attack from male courtship attempts at undesirable times or locations (Hews et al. 2004). Lastly, it is plausible that female aggression is an important component of courtship between the sexes, and a mechanism through which females can assess qualities of potential mates (Kralj-Fišer et al. 2013; DiRienzo et al. 2019). Female-male aggression has been reported in very few fly species, so *L. albimaculata* provides a useful system for investigating why such behaviours evolve.

### The hopping mud fly, *Lispe xenochaeta*

This species was only seen courting twice during the period of filming. Males are unique in that they perform side-ways ‘hops’ during courtship, which may act as a visual signal like the side-to-side dances exhibited by male *L. cana* (White et al. 2020). Also unique to this species is that while holding the females from behind, male *L. xenochaeta* vigorously ‘thump’ their mid-legs and ‘flick’ their wings, which almost certainly produces vibrational and aural cues, as in species of *Drosophila* (Fabre et al. 2012) and *Liriomyza* (Ge et al. 2018). This suggests that rather than solely as a form of mate guarding, ‘holding’ also serves to establish female receptivity in the lead-up to mating – this may also be true for *L. sydneyensis* and *L. cana*. Regarding the courtship behaviours of females, they seem to use the abdomen as a sexual signal, alternating between spread and closed wings to either hide or display their patterned abdomens. Most *Lispe* species have such patterned abdomens, and they are generally species-specific with differences in the shape and position of white markings (Pont 2019). It is possible that these abdominal patterns are involved as species-or sex-specific cues during courtship, as in many other invertebrates (Girard et al. 2011; Agrawal and Dickinson 2019). In one of the interactions, we observed that an unreceptive and aggressive female did not spread her wings to reveal her abdomen. As such, it seems plausible that the female ‘wing spread’, ‘wing quiver’, and abdomen display act as signals of female receptivity to the male. Importantly however, we only observed two interactions between males and females in this species, so there may be other inter-or intra-sexual interactions that occur. Regarding female aggression, similarly to *L. albimaculata*, female *L. xenochaeta* can be aggressive towards males – charging and attacking them during courtship events. This may be a response to male aggression and forced copulation attempts – whereby males repeatedly ‘tackle’ females during courtship. As mentioned above, there are also various other reasons that female-male aggression can occur, including female mate-assessment, or following the reception of male ejaculate, and *L. xenochaeta* provides ample opportunity for testing such hypotheses. Overall, these remarkable species further highlight the many behavioural complexities that are expressed by calyptrate flies during mating. Due to the ease with which they can be observed and collected, *Lispe* provide promising opportunities to investigate behavioural and evolutionary questions – and there is much to be gained from investigating the underpinnings of male-male competition and female-male aggression in the species highlighted here. Given that *Lispe* species can be easily found worldwide and exhibit wildly diverse behaviours, we encourage researchers to consider them as model species in their own studies of animal evolution, behaviour, and ecology.

## SUPPLEMENTARY MATERIAL

Supplementary Material 1 – Courtship behaviour of *Lispe sydneyensis*: https://youtu.be/rIAJY7p2ql0

Supplementary Material 2 – Courtship behaviour of *Lispe albimaculata*: https://youtu.be/k6BKLK4Dkwc

Supplementary Material 3 – Courtship behaviour of *Lispe xenochaeta*: https://youtu.be/gscgzAqqFYI

## Acknowledgements

The authors thank Adrian Pont for his helpful comments and guidance with taxonomic identification, Kathryn Doty for her assistance in the field, and Penelope Butterworth for her continuous support.

## REFERENCES

Agrawal S, Dickinson MH (2019) The effects of target contrast on Drosophila courtship. Journal of Experimental Biology 222:jeb203414

Arnqvist G, Henriksson S (1997) Sexual cannibalism in the fishing spider and a model for the evolution of sexual cannibalism based on genetic constraints. Evolutionary Ecology 11:255–273.

Bath E, Bowden S, Peters C, Reddy A, Tobias JA, Easton-Calabria E, Seddon N, Goodwin SF, Wigby S (2017) Sperm and sex peptide stimulate aggression in female Drosophila. Nature Ecology and Evolution 1:0154.

Bath E, Edmunds D, Norman J, Atkins C, Harper L, Rostant WG, Chapman T, Wigby S, Perry J (2021) Sex ratio and the evolution of aggression in fruit flies. Proceedings of the Royal Society B 299:20203053

Benelli G, Canale A, Bonsignori G, Ragni G, Stefanini C, Raspi A (2012) Male wing vibration in the mating behavior of the olive fruit fly Bactrocera oleae (Rossi) (Diptera: Tephritidae). Journal of Insect Behavior 25:590–603.

Bonduriansky R, Brooks RJ (1999) Why do male antler flies (Protophilia litigata) fight? The role of male combat in the structure of mating aggregations on moose antlers. Ethology Ecology and Evolution 11:287–301.

Butterworth NJ, Byrne PG, Wallman JF (2019) The blowfly waltz: Field and laboratory observations of novel and complex dipteran courtship behavior. Journal of Insect Behavior 32:109–119.

Butterworth NJ, White TE, Byrne PG, Wallman JF (2021) Love at first flight: Wing interference patterns are species-specific and sexually dimorphic in blowflies. Journal of Evolutionary Biology 34:558–570.

DiRienzo N, Bradley CT, Smith CA, Dornhaus A (2019) Bringing down the house: male widow spiders reduce the webs of aggressive females more. Behavioural Ecology and Sociobiology 73:1.

Fabre C, Hedwig B, Conduit G, Lawrence PA, Goodwin SF, Casal J (2012) Substrate-borne vibratory communication during courtship in Drosophila melanogaster. Current Biology 22:2180–2185

Fogaça JM, de Carvalho CJB (2018) Neotropical Lispe (Diptera: Muscidae): notes, redescriptions and key to species. Journal of Natural History 52:2147–2184.

Fordyce JA, Nice CC, Forister ML, Shapiro AM (2002) The significance of wing pattern diversity in the Lycaenidae: mate discrimination by two recently diverged species. Journal of Evolutionary Biology 15:871–879.

Frantsevich L, Gorb S (2006) Courtship dances in the flies of the genus Lispe (Diptera: Muscidae): from the fly’s viewpoint. Archives of Insect Biochemsitry 62:26–42.

Ge J, Wie J, Zhang D, Hu C, Zheng D, Kang L (2018) Pea leafminer Liriomyza huidobrensis (Diptera: Agromyzidae) uses vibrational duets for efficient sexual communication. Insect Science 26:510–522

Girard M, Kasumovic M, Elias D (2011) Multi-modal courtship in the peacock spider, Maratus volans (O.P.-Cambridge, 1874). PLOS ONE 6:e25390

Hews DK, Castellano M, Hara E (2004) Aggression in females is also lateralized: left-eye bias during aggressive courtship rejection in lizards. Animal Behaviour 68:1201–1207

Hohmann G, Fruth B (2003) Intra- and inter-sexual aggression by bonobos in the context of mating. Behaviour 140:1389–1413.

Kralj-Fišer S, Sanguino Mostajo GA, Preik O, Pekár S, Schneider JM (2013) Assortative mating by aggressiveness type in orb weaving spiders. Behavioral Ecology 24:824–831.

Kuramitsu K, Yooboon T, Tomatsuri M, Yamada H, Yokoi T (2019) First come, first served: precopulatory mate-guarding behavior and male-male contests by a hymenopteran saproxylic parasitoid. The Science of Nature 106:23.

Kutty SN, Pape T, Wiegmann BM, Meier R (2010) Molecular phylogeny of the Calyptratae (Diptera: Cyclorrhapha) with an emphasis on the superfamily Oestroidea and the position of Mystacinobiidae and McAlpine’s fly. Systematic Entomology 35:614–635

Land MF, Collett TS (1974) Chasing behaviour of houseflies (Fannia canicularis). Journal of Comparative Physiology 89:331–357.

Maklakov AA, Bilde T, Lubin Y (2004) Sexual selection for increased male body size and protandry in a spider. Animal Behaviour 68:1041–1048.

McAlpine DK, Kent DS (1981) Systematics of Tapeigaster (Diptera: Heleomyzidae) with notes on biology and larval morphology. Proceeding of the Linnean Society of New South Wales 106:33–58.

Morley EL, Steinmann T, Casas J, Robert D (2012) Directional cues in Drosophila melanogaster audition: structure of acoustic flow and inter-antennal velocity differences. Journal of Experimental Biology 215:2405–2413.

Parker GA (1970) The reproductive behaviour and the nature of sexual selection in Scatophaga stercoraria L. (Diptera: Scatophagidae). Behaviour 37:140–168.

Pont AC (2019) Studies on the Australian Muscidae (Diptera). VIII. The genus Lispe Latreille, 1797. Zootaxa 4557:001–232.

Pont AC, Meier R (2002) The Sepsidae (Diptera) of Europe. Fauna Entomologica scandinavica. Brill, Leiden, The Netherlands

Spofford M, Kurczewski F (1985) Courtship and mating behavior of Phrosinella aurifacies Downes (Diptera: Sarcophagidae: Miltogramminae). Proceedings of the Entomological Society of Washington 87:273–282.

Swierk L, Langkilde T (2013) Sizing-up the competition: Factors modulating male display behavior during mate competition. Ethology 119:948–959

Vikhrev N (2011) Review of the Palaearctic members of the Lispe tentaculate species-group (Diptera, Muscidae): revised key, synonymy and notes on ecology. Zookeys 84:59–70.

Werner D, Pont AC (2006) The feeding and reproductive behaviour of the Limnophorini (Diptera: Muscidae). Proceedings of the International Symposium of Simuliidae 14:79–114.

White TE, Vogel-Ghibely N, Butterworth NJ (2020a) Flies exploit predictable perspectives and backgrounds to enhance iridescent signal salience and mating success. The American Naturalist 195:733–742

White TE, Latty T (2020b) Flies improve the salience of iridescent sexual signals by orienting toward the sun. Behavioral Ecology 31:1401–1409

Zhang D, Wang Q, Liu X, Li K (2013) Sensilla on antenna and maxillary palp of predaceous fly, Lispe neimongola Tian et Ma (Diptera: Muscidae). Micron 49:33–39.

